# On the Mkv Model with Among-Character Rate Variation

**DOI:** 10.1101/2024.11.15.623796

**Authors:** Alessio Capobianco, Sebastian Höhna

## Abstract

Models used in likelihood-based morphological phylogenetics often adapt molecular phylogenetics models to the specificities of morphological data. Such is the case for the widely used Mkv model— which introduces an acquisition bias correction for sampling only characters that are observed to be variable—and for models of among-character rate variation (ACRV), routinely applied by researchers to relax the equal-rates assumption of Mkv. However, the interaction between variable character acquisition bias and ACRV has never been explored before. We demonstrate that there are two distinct approaches to condition the likelihood on variable characters when there is ACRV, and we call them joint and marginal acquisition bias. Far from being just a trivial mathematical detail, we show that the way in which the variable character conditional likelihood is calculated results in different assumptions about how rate variation is distributed in morphological datasets. Simulations demonstrate that tree length and amount of ACRV in the data are systematically biased when conditioning on variable characters differently from how the data was simulated. Moreover, an empirical case study with extant and extinct taxa reveals a potential impact not only on the estimation of branch lengths, but also of phylogenetic relationships. We recommend the use of the marginal acquisition bias approach for morphological datasets modeled with ACRV. Finally, we urge developers of phylogenetic software to clarify which acquisition bias correction is implemented for both estimation and simulation, and we discuss the implications of our findings on modeling variable characters for the future of morphological phylogenetics.

---

Discrete morphological characters have been used to reconstruct the evolutionary history of organisms since the dawn of phylogenetic systematics (Hennig, 1965; Kluge and Farris, 1969). Despite the molecular phylogenetic revolution providing an unprecedented amount of data and shaking up major portions of the Tree of Life (Savolainen and Chase, 2003; Delsuc et al., 2005; Near and Thacker, 2024), morphology remains the exclusive source of data to infer the phylogenetic relationships of extinct species—barred exceptional preservation of molecules in deep time (Hagelberg et al., 2015; Paterson et al., 2024). Moreover, interest in morphological phylogenetics is currently on the rise, thanks to methodological advances allowing for the joint estimation of phylogeny and divergence times of both extant and extinct taxa combined (Zhang et al., 2016; Wright et al., 2022), to improved morphological data collection and availability (Davies et al., 2017; Blackburn et al., 2024; Goswami and Clavel, 2024), and to the recognition that ignoring data from extinct species can bias inferences on macroevolutionary processes (Slater et al., 2012; Betancur-R et al., 2015; Lloyd and Slater, 2021; Faurby et al., 2024; Goswami and Clavel, 2024).

Likelihood-based methods for morphological phylogenetics have been introduced at the beginning of the 21^st^ century by adapting continuous-time Markov models developed for molecular data to the specifics of discrete morphological data (Lewis, 2001; Nylander et al., 2004). The core of likelihood-based morphological phylogenetics is the Mk model (Markov *k*-states model). Under the Mk model, all characters have the same expected number of changes across the tree—that is, all characters share the same rate of evolution. Moreover, for each character, each of the *k* states is equally probable at equilibrium. The combination of equal rates and equal state frequencies results in identical rates of transition between states (Lewis, 2001). When *k* =4, the Mk model is identical to the Jukes-Cantor model (Jukes et al., 1969) commonly used in molecular phylogenetics.

Several extensions to the Mk model have been introduced as an attempt to relax its strict assumptions, and to build models more appropriate to the nature of discrete morphological data. Firstly, Lewis (2001) already recognized a crucial distinction between the structure of discrete morphological datasets and that of molecular datasets. While molecular data matrices usually include every site—nucleotide or amino acid—between start point and end point of a sequence (no matter whether those sites are observed to be variable or invariant), morphological data matrices tend to not include characters that are not observed to vary among the sampled species. Branch lengths are systematically overestimated under the Mk model when calculated from only variable characters (Lewis, 2001).

To solve the problem of the absence of constant characters in discrete morphological data, Lewis (2001) proposed to compute a conditional likelihood, where the condition is to sample only characters that are observed to be variable. This is done by dividing the likelihood for a character by the probability to observe that character as variable. Effectively, the probability to observe a character as variable is more easily calculated as the complement of the probability to observe that character as constant or invariant (Lewis, 2001; see Fig. 1 for how the terms invariant and variable character are used in this paper). The model that uses this conditional likelihood was termed Mkv—where the ‘v’ stands for ‘variable’—to distinguish it from the Mk model (Lewis, 2001). The Mkv model is also sometimes referred to as the Mk model with acquisition bias correction (or ascertainment bias correction; e.g., Lam et al., 2018, Matzke and Irmis, 2018).

**Figure 1:**
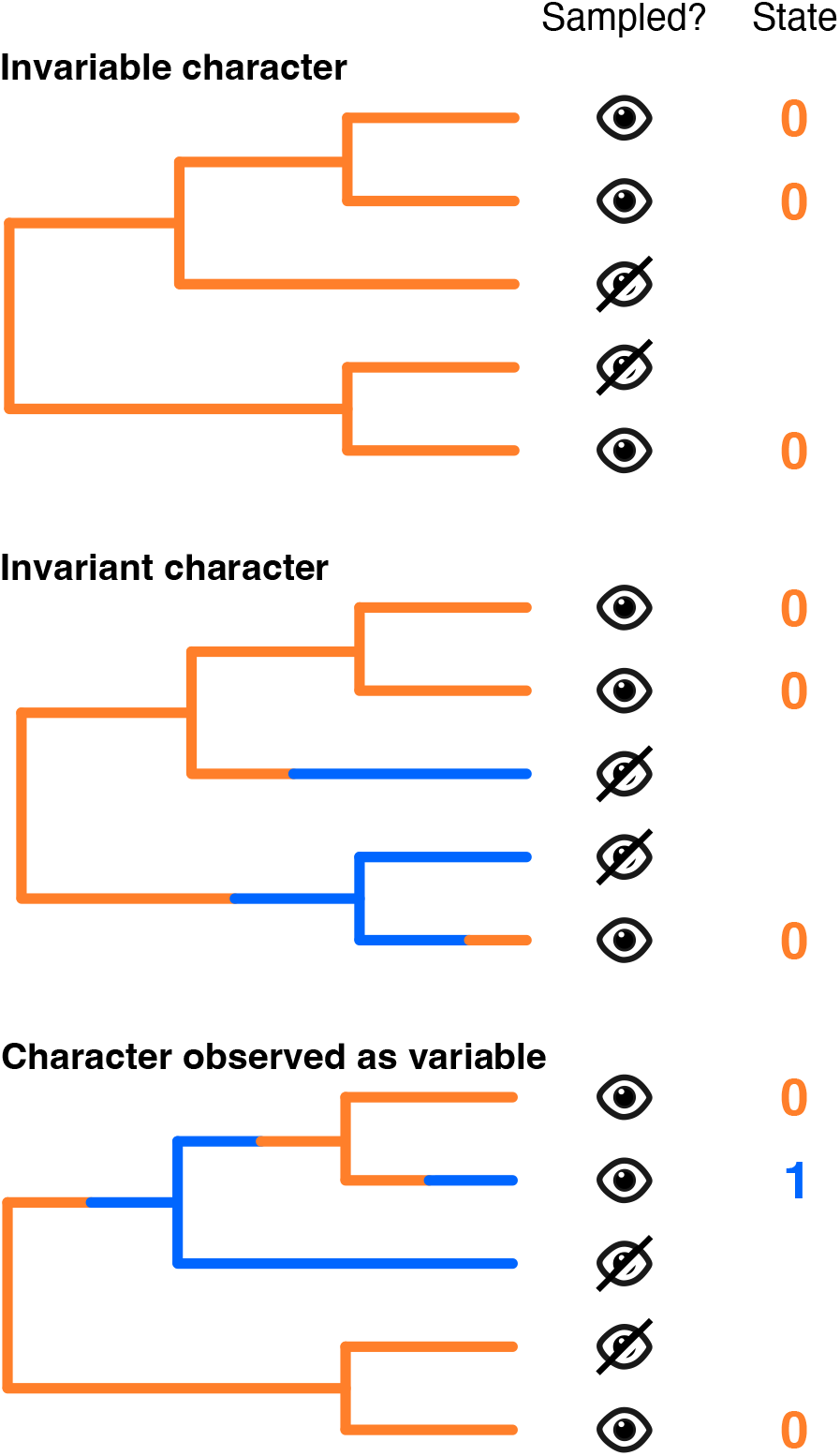
Differences between invariable character, invariant character, and character observed as variable. Invariable characters are a subset of invariant characters that are always constant (their state space is effectively equal to 1). For simplicity, we sometimes use the term ‘variable character’ in the text to indicate a character that is observed to be variable. The Mkv model assumes that all sampled characters are observed as variable.

Another major extension to the Mk model comes with relaxing the assumption of equal rates of evolution for different characters, i.e., allowing for among-character rate variation (ACRV). This is done by modeling ACRV with a flexible distribution that can accommodate from (almost) no rate variation up to high amounts of rate variation, depending on the value of an estimated parameter that determines the shape of that distribution. The most common distribution used is the discretized gamma distribution, which approximates a continuous gamma distribution with a discrete number of equiprobable categories, representing mean (or median) points between quantiles of that distribution (Yang, 1994). In molecular phylogenetics, 4 discrete categories are often applied, as originally recommended by Yang (1994) based on likelihood values plateauing above 4 categories for different molecular datasets. However, a larger number of categories (7 to 14) might be more appropriate for morphological data (Wright and Wynd, 2024). Alternatively, a discretized lognormal distribution is sometimes used to model ACRV in morphological datasets, as some studies have shown that a lognormal distribution might be a better fit in datasets where most characters have a relatively low rate (a condition expected under atomistic character coding, where complex phenotypes are broken into smallest discernible character units; Wagner, 2012; Harrison and Larsson, 2015). Theoretically, rates could be sampled directly from the full continuous distribution (whether gamma, or lognormal, or other) without discretizing it in a set number of rate categories (Yang, 1993), but this is rarely—if ever—done empirically.

Every major likelihood-based phylogenetic software allows to apply the Mkv model together with a model of ACRV when analyzing a morphological dataset (e.g., `RevBayes`, `MrBayes`, `BEAST2`, `IQ-TREE`, `RAxML`; Ronquist et al., 2012; Bouckaert et al., 2014; Stamatakis, 2014; Höhna et al., 2016; Minh et al., 2020), and this is done routinely in empirical studies. While Harrison and Larsson (2015) pointed out that unsampled constant characters are more probable under lower rate categories, potentially leading to larger corrections in ACRV models relative to an equal-rates model (Harrison and Larsson, 2015), the effects of variable character acquisition bias on ACRV analyses remain surprisingly unexplored. Moreover, how the acquisition bias correction should be implemented in cases of rate heterogeneity has not been explicitly discussed in the literature to our knowledge.

In this paper, we show two distinct ways—both mathematically viable—to condition the likelihood on variable characters when there is more than one rate category. We explore the implications that these two different conditional likelihood approaches have regarding how rate variation is distributed in morphological datasets, and show how they impact phylogenetic analyses using both simulated and empirical data. Finally, we provide insights into future model development in morphological phylogenetics—such as continuous distributions for ACRV and models with unequal state frequencies—in light of what we learned about the application of variable character acquisition bias to rate-heterogeneous models.

## 1 Modeling Acquisition Bias with ACRV

In the Mkv model described by Lewis (2001), the tree likelihood for character *i* conditional on character *i* being observed as variable is calculated as follows:

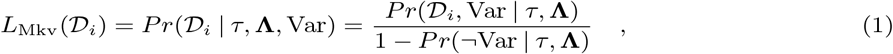

where *𝒟*_*i*_ is the observed data associated with character *i, τ* is the tree topology, **Λ** is the vector of branch lengths *λ*_1_, *λ*_2_, *λ*_3_, …, and ‘Var’ and ‘¬Var’ refer to observing character *i* respectively as variable or invariant (see also Fig. 1). The probability of observing a character to be invariant is:

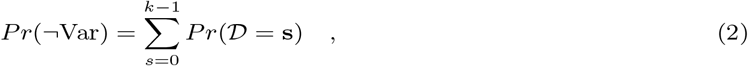

where *k* is the number of states, and *Pr*(*𝒟* = **s**) is the probability of character states being equal to *s* for all observed samples of one character. These are obtained by calculating the likelihoods for *k* dummy characters having the same state for every tip of the tree (one having all ‘0’s, one having all ‘1’s, and so on). If transition rates and state frequencies at equilibrium are equal (as in the Mkv model), then *Pr*(*𝒟* = **0**) = *Pr*(*𝒟* = **1**) = … = *Pr*(*𝒟* = **k** − **1**), and Equation 2 becomes:

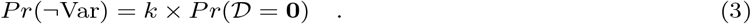

If branch lengths are the same for every character (i.e., there is only one rate category) as it is the case in the Mkv model, then the likelihood for a character matrix with *n* characters under the Mkv model is:

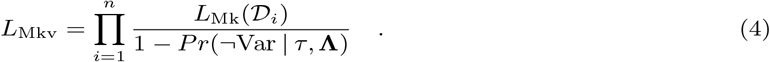

We will now show that the variable character conditional likelihood can be calculated in two different ways when there is more than one rate category. From now on and in the following sections, we will take into consideration an ACRV model with a discretized gamma distribution, but the same likelihood equations and general concepts are also valid for a discretized lognormal ACRV model—or indeed any ACRV model with multiple rate categories.

Under a discretized gamma model with *r* rate categories *ρ*_1_, *ρ*_2_, …, *ρ*_*r*_, each of them with probability *Pr*(*ρ*_*j*_), the likelihood for a character matrix with *n* characters *without* variable character acquisition bias correction is equal to:

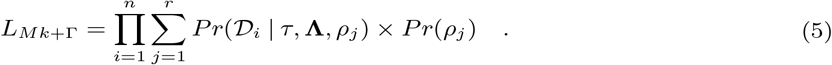

Here, rate categories *ρ*_1_, *ρ*_2_, …, *ρ*_*r*_ act as multipliers to the vector of branch lengths **Λ**. In order for the model to be identifiable, the mean of rate categories has to be equal to 1 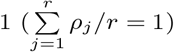.

One approach to calculate the variable character conditional likelihood under the discretized gamma model (or any other model with multiple rate categories) is to condition the likelihood *by rate category*. We call this *joint acquisition bias* correction, and the resulting model jMkv + Γ. The likelihood for this model is calculated as follows:

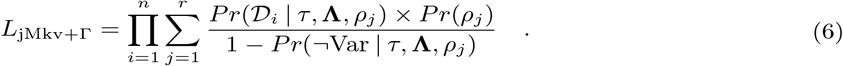

Under the jMkv + Γ model, for each rate category the likelihood of character *i* under that rate is divided by the probability that character *i* is observed to be variable given that rate.

The other approach to calculate the variable character conditional likelihood under the discretized gamma model (or any other model with multiple rate categories) is to condition the likelihood *over all rate categories*. We call this *marginal acquisition bias* correction, and the resulting model mMkv + Γ. The likelihood for this model is calculated as follows:

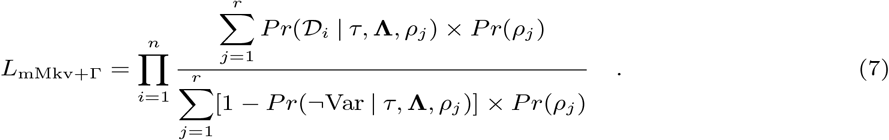

Under the mMkv + Γ model, the likelihood of character *i* integrated over all rate categories is divided by the summation of probabilities of character *i* being observed to be variable under each rate.

Notice that these two approaches (joint acquisition bias and marginal acquisition bias corrections) are identical and collapse into the acquisition bias correction of Lewis’ Mkv model when there is only one rate category (*r* = 1). Thus, both jMkv and mMkv are mathematically consistent with the Mkv model.

## 2 Simulating Data under Acquisition Bias with ACRV

The effective difference between joint acquisition bias correction (jMkv model) and marginal acquisition bias correction (mMkv model) can be better understood by looking at how data are simulated under these two approaches. Under the joint acquisition bias approach, the correction is larger for lower rate categories and smaller for higher rate categories, because it is more likely that lower rates would generate invariant characters (as pointed out by Harrison and Larsson, 2015). From a simulating perspective, this corresponds to first drawing a rate category from the prior distribution, then simulating a character under that rate until the character is observed to be variable (Fig. 2a). This process is repeated for how many characters compose the simulated data matrix.

**Figure 2:**
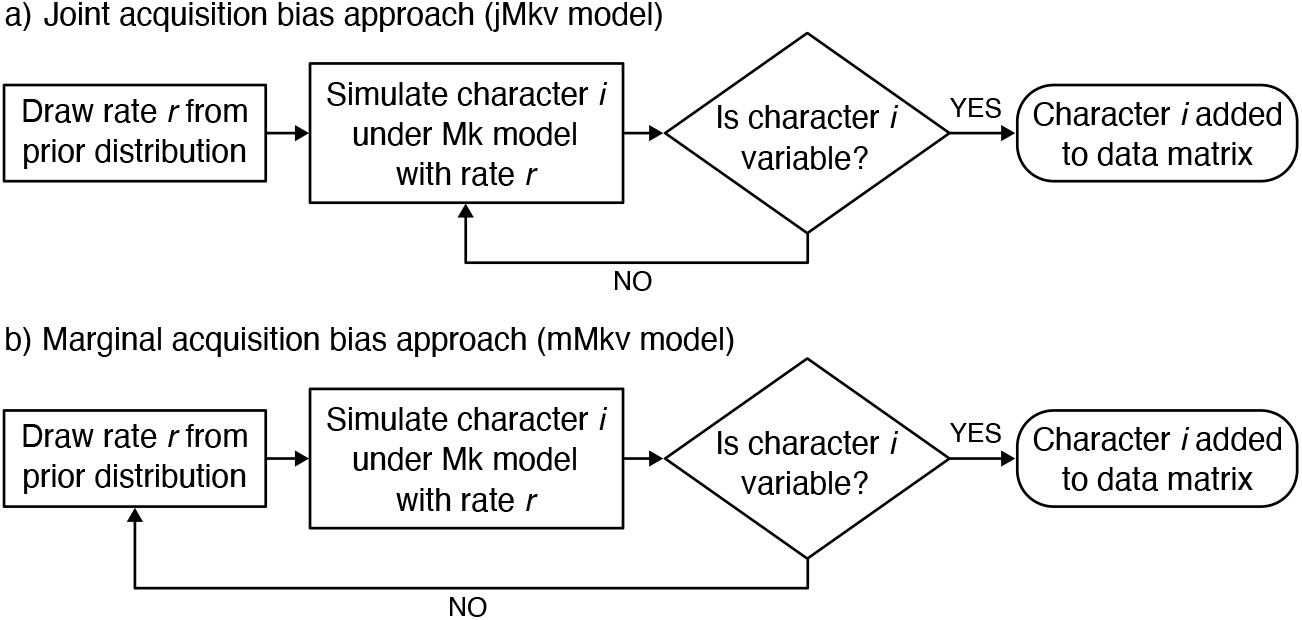
Schematic graph of the two approaches to simulate data with ascertainment bias. The outlined steps are repeated for each character. a) Joint acquisition bias approach, corresponding to the jMkv model. b) Marginal acquisition bias approach, corresponding to the mMkv model.

The marginal acquisition bias approach does not correct the tree likelihood per rate category. Instead, it calculates a marginal correction that integrates over all rate categories, and applies it to the overall tree likelihood computed as it would have no acquisition bias. From a simulating perspective, instead of simulating a character under the same rate until that character is variable, the rate category is redrawn from the prior distribution every time the simulated character is not observed to be variable (Fig. 2b).

We wrote an `R` script to generate data matrices under the jMkv+Γ and mMkv+Γ models, given a phylogeny (topology + branch lengths). The logical steps for simulating the data follow those outlined in Fig. 2. An unrooted phylogeny of 20 terminal nodes (tips) was simulated in the software RevBayes (Höhna et al., 2016), using a uniform prior on topology and an exponential prior with mean = 0.1 (*λ* = 10) on each branch length. The gamma distribution modeling ACRV was set to have an alpha parameter *α* = 1, and it was discretized in four rate categories. The function `discrete.gamma` in the `R` package `phangorn` (Schliep, 2011) was used to get the values of rate categories. The function `sim.Mk` in the `R` package `phytools` (Revell, 2012) was used to simulate each character under an Mk model with a rate drawn from the four equiprobable rate categories (Fig. 2). A matrix of 10,000 binary characters was generated under both the jMkv+Γ and mMkv+Γ models, and for each character the rate under which it was generated was also stored in a separate text file. To explore the effect of tree length in the distribution of per-character rates, the simulation was repeated for a rescaled phylogeny, where branch lengths of the original generating phylogeny were rescaled by a factor of 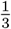.

Additionally, we also generated data matrices under jMkv and mMkv models where the per-character rates are drawn directly from a continuous gamma distribution, rather than from its discretized version. This was done to show more clearly the relationship between the rate distribution from which rates are drawn and the effective distribution of rates in the character matrix under both types of acquisition bias correction. The steps to simulate data were kept identical to the ones listed above, except for the use of a continuous gamma distribution instead of a discretized one.

Fig. 3 shows the effective distributions of rates of character evolution under the jMkv+Γ, mMkv+Γ, jMkv + continuous gamma, and mMkv + continuous gamma models. Under jMkv, the effective distribution of evolutionary rates for the observed (variable) characters follows the underlying ACRV distribution. In other words, if ACRV is modeled with a 4-categories discretized gamma distribution where each rate category is equiprobable, then roughly 25% of observed variable characters has evolved under the lowest rate, and 25% under the highest rate (Fig. 3). As a corollary, the relative rates of observed characters are independent from the tree length. That is, the distribution of per-character relative rates will be the same no matter what the average number of state transitions across the whole tree is.

**Figure 3:**
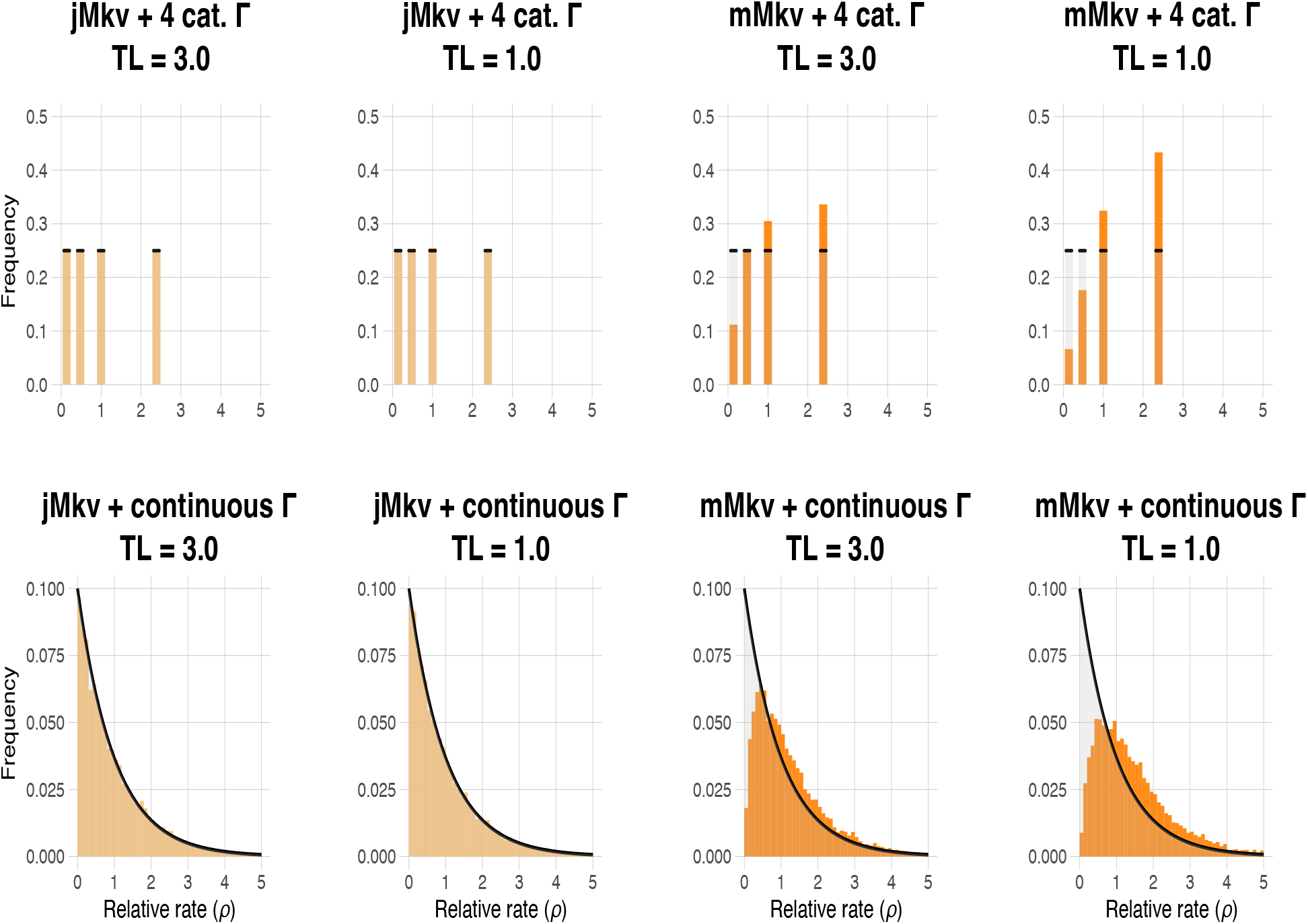
Effective distributions of evolutionary rates when 10,000 characters are simulated under the two variable character acquisition bias approaches. Top row: models with 4-categories discretized gamma distribution of ACRV; bottom row: models with continuous gamma distribution of ACRV. In the top row, each bar represents a rate category, and the black ticks mark the expected frequency of each category under no acquisition bias. In the bottom row, the probability density for a continuous gamma distribution with alpha = 1 is shown in the background (rescaled by 0.1). Notice how, under the mMkv model, slower rates are underrepresented among observed variable characters, and the effective rate distribution is right-skewed compared to the true rate distribution that generates the characters. TL = tree length.

Conversely, under mMkv the effective distribution of evolutionary rates for the observed (variable) characters is right-skewed compared to the underlying ACRV distribution. If ACRV is modeled with a 4-categories discretized gamma distribution where each rate category is equiprobable, then the lowest rate category will be underrepresented among the observed variable characters, whereas the highest rate category will be overrepresented. This effect is stronger the shorter the tree is (Fig. 3). Thus, under mMkv the mean evolutionary rate of observed variable characters is higher than the mean rate of the underlying distribution of rates under which characters were generated.

The marginal acquisition bias approach converges to the joint acquisition bias approach as the tree length increases tending towards infinity. Longer tree lengths imply that characters simulated under the lowest rate category will be more often observed to be variable. Thus, simulated characters will always be variable when tree length approaches infinity, and the effective rate distribution under mMkv will be equal to the effective rate distribution under jMkv (and to the underlying ACRV distribution).

## 3 Simulation Study

We designed a simulation study to explore the performance of the two variable character acquisition bias approaches (joint and marginal) under different scenarios. Simulated data composed of 1,000 binary characters for 20 operational taxonomic units (OTUs) were generated under the following three procedures:

1. No acquisition bias. Each character is simulated under a rate drawn from a discretized gamma distribution. Both variable and invariant characters are kept in the character matrix. This corresponds to a Mk+Γ model.
2. Joint acquisition bias (Fig. 2a). After a rate is drawn from a discretized gamma distribution, a character is simulated until it is observed to be variable. Then it is added to the character matrix, which is made of variable characters only. This corresponds to a jMkv+Γ model.
3. Marginal acquisition bias (Fig. 2b). A character is simulated under a rate drawn from a discretized gamma distribution. As long as the character is not observed to be variable, a new rate is drawn from the same distribution, and the character is simulated again. When the character is observed to be variable, it is added to the character matrix, which is made of variable characters only. This corresponds to a mMkv+Γ model.

Character evolution was simulated on an unrooted phylogeny of 20 tips, which was generated in `RevBayes` using a uniform prior on topology and an exponential prior with mean = 0.1 (*λ* = 10) on each branch length. As in the previous section, the gamma distribution modeling ACRV was set to have an alpha parameter = 1 and discretized in four rate categories. Characters were simulated in `R` using the same procedure outlined in the previous section. The simulations were replicated on 250 different phylogenies, each generated under the same priors. Overall, we simulated 750 datasets (3 simulation scenarios *𝒟* 250 phylogenies), each with 1,000 binary characters per 20 OTUs.

We analyzed each simulated dataset under 5 different substitution models:

1. Mk (no acquisition bias correction; no ACRV);
2. Mkv (acquisition bias correction; no ACRV);
3. Mk+Γ (no acquisition bias correction; gamma-distributed ACRV);
4. jMkv+Γ (joint acquisition bias correction; gamma-distributed ACRV);
5. mMkv+Γ (marginal acquisition bias correction; gamma-distributed ACRV).

For all the models with ACRV, the gamma distribution modeling ACRV was discretized into 4 categories. The prior on the alpha parameter determining the shape of the gamma distribution was set as uniform between 0 and 10^8^, following the recommendation of Fabreti and Höhna (2023). The priors on the phylogeny were set as an unrooted uniform prior on topology and an exponential prior with mean = 0.1 (*λ* = 10) independently on each branch length. When applying the models with acquisition bias correction (Mkv, jMkv+Γ, and mMkv+Γ), invariant characters were pruned from the datasets simulated under no acquisition bias (Mk+Γ model). This is because including invariant characters biases the likelihood calculated with acquisition bias (Leaché et al., 2015; see also our *Discussion*). For each analysis, four independent MCMC (Markov chain Monte Carlo) simulations were run for 50,000 iterations, sampling trees and parameters every 10 iterations. The first 10% of each MCMC simulation was discarded as burn-in. All analyses were run in RevBayes v.1.2.2 (Höhna et al., 2016) on the PalMuc high-performance computing cluster (HPC) at LMU Munich.

Convergence of continuous parameters and tree parameters between independent replicates for each simulated data analysis was assessed using the `R` package `convenience` (Fabreti and Höhna, 2022). The threshold for minimum effective sample size (ESS) of continuous parameters was set as 200. Convergence of continuous parameters between independent runs was assessed with the Kolmogorov-Smirnov (KS) test, with the KS value threshold calculated at precision level = 0.0177 (Fabreti and Höhna, 2022). The threshold for maximum difference in split frequencies of phylogenies corresponded to the expected difference in split frequencies calculated for ESS = 200 (precision = 0.0177) (Fabreti and Höhna, 2022). Analyses that did not reach convergence according to this protocol were run again for 100,000 iterations, and their convergence reassessed.

### 3.1 Results of Simulation Study

Our simulation study shows that the choice of variable character acquisition bias approach has a substantial impact on the estimates of tree length and amount of ACRV (Fig. 4).

**Figure 4:**
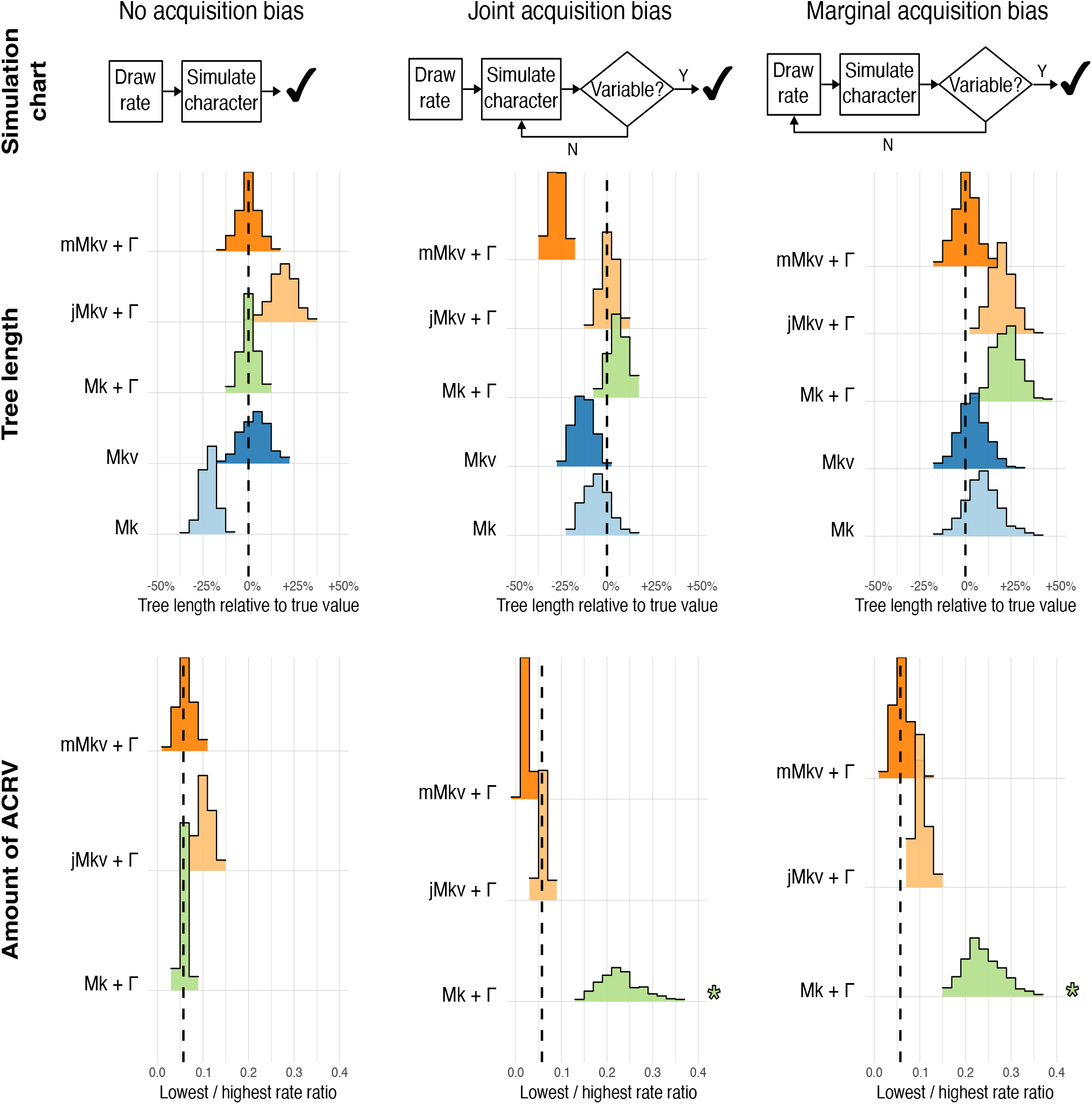
Bar plots of estimated mean tree length (showed as percentage difference relative to true value) and amount of among-character rate variation (showed as ratio between lowest and highest rate) inferred under different morphological substitution models for 250 simulated datasets three different simulation scenarios. Dashed lines indicate true values under which the data were simulated. Bar widths are equal to 5% for the top row (tree length) and to 0.02 for the bottom row (amount of ACRV). Some simulated datasets (7 simulated under joint acquisition bias and 5 under marginal acquisition bias) were estimated to have no ACRV (lowest-to-highest rate ratio *>* 0.99) under Mk+Γ; these have been removed from the bar plot and signaled by an asterisk.

When both variable and invariant characters are simulated (under the Mk+Γ model), the mMkv+Γ model clearly outperforms the jMkv+Γ model in correctly estimating both tree length and amount of ACRV. This is consistent with how the marginal acquisition bias approach works (Fig. 3), as it assumes that the observed variable characters evolve under a higher average rate than the true rate of evolution for all characters (variable + invariant). Conversely, the jMkv+Γ model assumes that the rate distribution of the observed variable characters is equal to the rate distribution among every character generated from the same process, including the invariant ones. This leads to an overestimation of the average rate of character evolution—and thus of the tree length—if lower rates are overrepresented among invariant characters and underrepresented among variable characters.

When data were simulated under either joint or marginal acquisition bias, the corresponding model (respectively jMkv+Γ and mMkv+Γ) performs the best in estimating tree length and ACRV (Fig. 4). Not accounting for acquisition bias when analyzing datasets with only variable characters recovers similar tree lengths to the jMkv+Γ model. However, it has a dramatic effect on the ACRV estimates, as it greatly underestimates the amount of ACRV in the dataset (overestimation of the alpha parameter). This effect can be so pronounced that in few simulated datasets (7 datasets simulated under jMkv+Γ and 5 datasets simulated under mMkv+Γ) alpha was estimated to be larger than 10^7^ under Mk+Γ, effectively corresponding to equal rates of evolution among characters (Fig. 4).

Our simulation study also shows that variable character acquisition bias and ACRV interact in complex ways, such that it is difficult to disentangle the effects of one or the other model component on tree length estimates. For example, while adding ACRV to an Mk model has the effect of reconstructing longer trees, applying the marginal acquisition bias correction shortens the reconstructed tree so much that the mMkv+Γ model estimates shorter trees on average than the Mkv model.

## 4 Empirical Example

To explore whether different approaches to model the variable character acquisition bias have a practical impact on empirical phylogenetic analyses, we picked the morphological dataset of Villa (2023) focused on Gekkota (geckos and close relatives) as an empirical case study. This dataset was chosen because of a few desired features:

- Dataset size: not too few but not too many species (30), and a relatively large number of morphological characters (846).
- Mix of extant and extinct (fossil) species.
- Large number of characters that are invariant among observed species (476, more than 56% of total sampled characters). This is because the matrix of Villa (2023) uses the character list of Conrad (2018) (modified by Villa et al., 2022), which is focused on a much broader clade (Squamata) than Gekkota. Thus, several characters that are variable among squamates are actually invariant within Gekkota. This feature of the dataset allows to compare models without acquisition bias correction and models that implement acquisition bias correction.

We analyzed the Gekkota dataset under the same substitution models we used to analyze the simulated data: Mk, Mkv, Mk+Γ, jMkv+Γ, and mMkv+Γ. Priors on gamma distribution and phylogeny were set as in the simulated data analyses. Invariant or ambiguous characters for all taxa were automatically pruned from the Gekkota dataset when applying the models with acquisition bias correction. For each analysis, four independent MCMC (Markov chain Monte Carlo) were run for 100,000 iterations, sampling tree and parameters every 10 iterations. The first 10% of each MCMC chain was discarded as burn-in. We repeated all the analyses above for a pruned Gekkota dataset where we excluded extinct species (fossil data). This was done to explore how different taxonomic sampling strategies might impact the performance of character evolution models. All analyses were run on `RevBayes` v.1.2.2 (Höhna et al., 2016) on the PalMuc high-performance computing cluster (HPC) at LMU Munich. Convergence of continuous parameters and tree parameters between independent replicates was assessed using the `R` package `convenience` (Fabreti and Höhna, 2022), using the same protocol explained for the simulation study.

To check whether different substitution models reconstruct different posterior distributions of tree topologies, we applied the same expected difference of split frequencies test that we used to asses topology convergence (Fabreti and Höhna, 2022). If all topology splits compared between two models fall within the threshold for maximum expected difference in split frequencies due to sampling stochasticity, then it means that the same posterior distribution of tree topologies was sampled under the two models. However, if some topology splits compared between two models fall outside the threshold, then it means that different topology distributions were sampled under the two models.

### 4.1 Results of Empirical Example

Estimated tree lengths for the Gekkota empirical dataset were impacted by model choice consistently with what we observed in the simulation study. Overall, longer trees were recovered when the model did not include an acquisition bias correction (Fig. 5). However, among models that do allow for ACRV, jMkv+Γ estimated trees that are about as long (for the full dataset) or longer (for the only extant dataset) than trees that were estimated with the Mk+Γ when including invariant characters, whereas mMkv+Γ estimated much shorter trees. Both ACRV models with a variable character acquisition bias correction (jMkv and mMkv) recovered similarly low values of the alpha parameter of the gamma distribution, indicating high levels of ACRV in the Gekkota dataset. These estimates contrast with those of the Mk+Γ model, which recovered higher values of the alpha parameter, indicating little ACRV in the full Gekkota dataset, and even no ACRV—i.e., rate homogeneity—in the only extant Gekkota dataset (Fig. 5).

**Figure 5:**
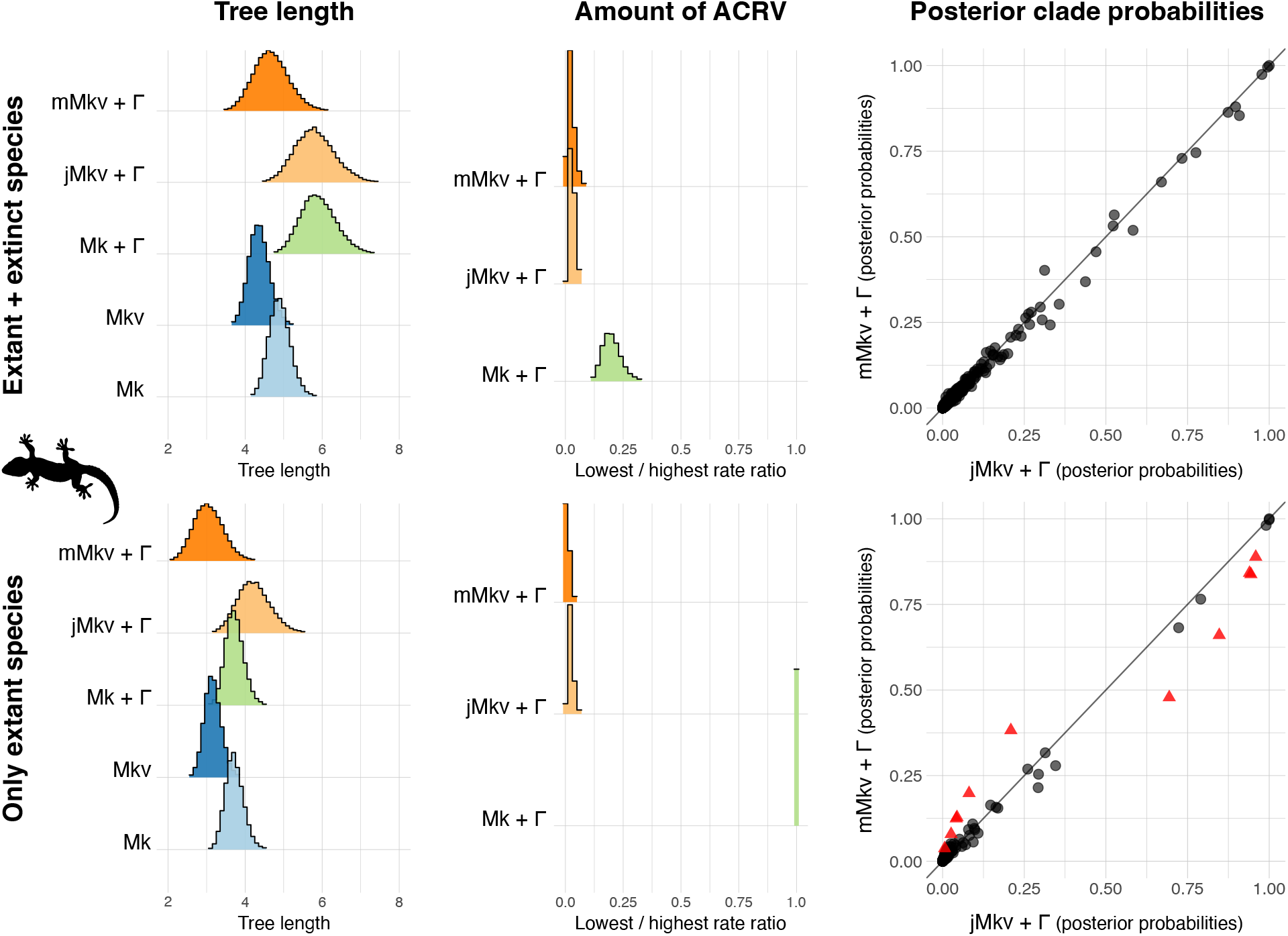
Impact of different models of morphological character evolution on estimated tree length (*left*), among-character rate variation (*center*; showed as ratio between lowest and highest rate), and tree topology (*right*) for the empirical Gekkota dataset, including (*top*) and excluding (*bottom*) extinct species. Tree lengths are expressed in average expected number of character transitions (rates × time). Bar widths are equal to 0.1 for the tree length plots, and to 0.02 for the rate ratio plots. Each symbol in the scatter plots represents one tree split (or clade). Circles indicate clades with posterior probabilities not significantly different under the expected difference of split frequencies test. Triangles indicate clades with significantly different posterior probabilities under the same test. Silhouette of *Gekko gecko* from PhyloPic (https://www.phylopic.org/).

The choice of substitution model not only affected continuous parameters such as tree length and shape of the gamma-distributed ACRV, but had also an effect on tree topology. Allowing for ACRV or not had the biggest impact on the posterior probabilities of clades (Supplementary Figs. 1 and 2). The type of variable character acquisition bias used when allowing for ACRV had a negligible impact on topology for the full Gekkota dataset, but had a significant impact for the only extant Gekkota dataset (Fig. 5). Specifically, the joint acquisition bias correction approach recovered higher probabilities for some clades with posterior probabilities *>* 0.45, while the marginal acquisition bias approach recovered higher probabilities for some low-probability clades (posterior probability *<* 0.45) (Fig. 5).

## 5 Discussion

### 5.1 Is an Acquisition Bias Correction Always Necessary for Discrete Morphological Datasets?

In this paper, we demonstrated that there are two different ways—both mathematically correct and consistent—to condition the tree likelihood on observing only variable characters when there is more than one rate of character evolution (rate heterogeneity). It might be tempting to consider variable character acquisition bias as a minor topic for the development of phylogenetic models of morphological evolution. After all, some morphological datasets—including the one used for our empirical case study— comprise invariant characters, so why should we even use models that condition on observing only variable characters? If a morphological character matrix includes also invariant characters, would it be more appropriate to use the Mk model (combined with a model of ACRV) rather than Mkv and its rate-heterogeneous derivations (jMkv and mMkv)?

The reason why invariant characters in morphological datasets cannot be modeled in the same way as in molecular datasets ultimately stems from ontological differences between molecular and morphological characters, rather than from researchers’ practices. Molecular characters are naturally discrete elements (nucleotides or amino acids) arranged in a linear fashion, whereas morphological characters often attempt to discretize an underlying continuous variation (Stevens, 1991; Wiens, 2001) and—more relevantly—are not linearly ordered. In molecular phylogenetics, after deciding a start point and an end point in a linear sequence (DNA or protein), every single character between those two points can be sampled— no matter whether it is variable or not among sampled taxa, how much it is variable, and whether an invariant character is invariant only within the studied clade or among a broader set of taxa. In morphological phylogenetics, even if a researcher would attempt to code more invariant morphological characters in their matrix, it is not clear at what point they should stop sampling invariant features to get an unbiased set of characters. For example, if they are building a matrix of cranial characters in apes, they could decide to include features that are invariant among apes but variable among primates. But they could also include features that are invariant among primates but variable among placentals; and those invariant among placentals but variable among mammals; and those invariant among mammals but variable among amniotes; and so on in a series of taxonomic Russian dolls. This means that a specific morphological dataset can have an arbitrarily varying number of invariant characters, which will result in differently estimated parameters under a model without an acquisition bias correction (such as the Mk model). Ultimately, it is impossible to unambiguously determine the number of invariant morphological characters for any set of sampled taxa.

Thus, we recommend to always use a model that conditions on observed characters being variable when analyzing discrete morphological datasets. In cases where the dataset also includes characters that are invariant or ambiguous among all sampled taxa, those characters should always be removed before the phylogenetic analysis—no matter if there are relatively few or many of them compared to the variable characters (see also Leaché et al., 2015 for why invariant characters need to be removed when applying the Mkv model).

### 5.2 How Should Acquisition Bias be Modeled for Rate-Heterogeneous Datasets?

Given the importance of the variable character acquisition bias correction for morphological phylogenetics, and the fact that it has been implemented in phylogenetic software allowing for rate heterogeneity in character evolution since two decades ago, it is perhaps surprising that—to our knowledge—the way in which the variable character acquisition bias is calculated when combined with ACRV has never been explicitly discussed. It is more likely that existing software implement what we called the mMkv model (or marginal acquisition bias correction), because it is more immediate to just divide the ‘usual’ tree likelihood computed without acquisition bias by a marginal correction integrated over rate categories. We checked the software `RevBayes`, `MrBayes`, and `BEAST2`, and they do indeed calculate a marginal acquisition bias correction when combining discretized gamma ACRV and acquisition bias correction for tree estimation. However, we urge software developers to indicate explicitly which acquisition bias correction (joint or marginal) is applied in their software, and to make sure that it is consistent between simulation and estimation.

The choice between mMkv and jMkv model ultimately stems from assumptions about how the evolutionary rates of observed variable characters are distributed compared to the rates of invariant characters. If the rate variation among morphological characters in the studied system—whether variable or invariant—is thought to be gamma-distributed (or lognormally-distributed; see Wagner, 2012 and Harrison and Larsson, 2015), and the characters that are observed to be identical are simply characters that evolve at a low rate and might be observed as variable with an increased taxonomic sampling (or given enough evolutionary time), then the mMkv model with the marginal acquisition bias correction is the appropriate model to choose.

The jMkv model assumes that characters observed to be variable and those observed as invariant do not share a common mechanism of evolution. In a sense, under the jMkv model every character that can potentially vary is observed to be variable. This is incredibly unlikely, as most often invariant morphological characters in empirical datasets are simply variable characters with a limited taxonomic sampling—as seen also in our empirical example. On such grounds, it is very difficult to justify the use of the jMkv model.

Besides theoretical arguments, our simulation study demonstrates that mMkv is accurate when all characters evolve under an Mk model with ACRV and the invariant ones are not sampled, while jMkv systematically overestimates tree length and underestimates rate heterogeneity under the same conditions (Fig. 4). In other words, mMkv manages to recover true tree length and amount of ACRV without the need to know the number of invariant characters—something that is impossible to know unequivocally for empirical morphological datasets, as we explained in the previous subsection. Thus, we recommend to use the marginal acquisition bias correction (mMkv model) to model variable characters together with rate heterogeneity.

#### 5.2.1 Does it matter?—

Our simulation study and empirical example show that the choice of acquisition bias approach will always have an impact on reconstructed tree length and—to a lesser extent—on the estimated amount of ACRV when there is rate variation in the dataset (this difference will disappear if there is no evidence of rate variation in the dataset, as both jMkv and mMkv will collapse into the Mkv model). In fact, the results we showed here help to explain some puzzling behaviour observed in previous morphological phylogenetics studies. For example, Mulvey et al. (2024) applied different models of morphological evolution to a broad set of empirical datasets, and found that adding a gamma-distributed ACRV to the Mk model slightly increased reconstructed tree length, while adding the same gamma-distributed ACRV to the Mkv model tended to greatly decrease reconstructed tree length. This result—while apparently confusing—is perfectly consistent with our explanation of how the marginal acquisition bias is calculated, and what assumptions it makes on the rate distribution of observed variable characters.

Moreover, we also showed through an empirical example that the choice of acquisition bias approach can have an impact on tree topology, although this will be dataset-dependent.

### 5.3 Implications for Models with Continuous ACRV

The existence of two viable but distinct ways to condition on observing variable characters with rate heterogeneity has to be taken into consideration for future model development in morphological phylogenetics. One potential development involves modeling ACRV under a continuous distribution rather than its discretized version. As mentioned previously, this has not been done empirically, despite the existence of a theoretical background (Yang, 1993) and available computational machinery. While the number of characters that are now included in standard molecular datasets (from tens of thousands to millions of characters) might pose a computational challenge for parameter-rich models such as continuous ACRV models, morphological matrices usually are much smaller, ranging from dozens to few thousand characters. Thus, continuous ACRV models might be feasible for morphological phylogenetics, and might allow for more fine-grained estimation of per-character evolutionary rates in morphological datasets.

For an ACRV model with a continuous distribution of rates, the tree likelihoods of the jMkv and mMkv models become, respectively:

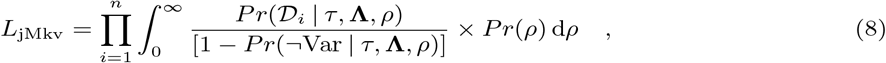

and:

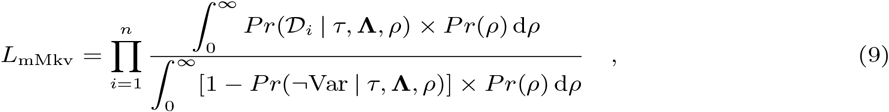

where *ρ* is the per-character relative rate, and *Pr*(*ρ*) is the prior distribution of *ρ* with domain between 0 and ∞. It is straightforward to implement the joint acquisition bias correction with continuous ACRV in existing Bayesian software machinery, as the denominator in Equation 8 can be calculated at each step of a Bayesian MCMC by calculating the likelihood of an invariant ‘dummy character’ given the parameters at that step. Furthermore, the integral in Equation 8 can be omitted and is solved instead with standard MCMC sampling (e.g., data augmentation by the rate category per character).

On the contrary, implementing the marginal acquisition bias correction is not immediate, because the integral at the denominator in Equation 9 cannot be solved through MCMC integration and has to be calculated through an approximate numerical solution (i.e., in the end using discretization to compute the integral). It will not be possible to compare the performance of continuous and discretized ACRV models with morphological data until ways to calculate the marginal acquisition bias correction are implemented in phylogenetic software. While this is not a trivial task, it would open the door for the development of more complex models in morphological phylogenetics, where each character is allowed to evolve at a different rate.

### 5.4 Implications for Models with Base-Composition Heterogeneity

All models of character evolution discussed in this paper assume that state frequencies for each character are equal at equilibrium. This is usually not the case in molecular phylogenetics, where state frequencies of nucleotides or amino acids are estimated as model parameters or empirically calculated based on their proportions in the data. Attempts to similarly model unequal state frequencies in morphological phylogenetics are greatly complicated by the arbitrary labeling of character states in morphological datasets, combined with the lack of a common meaning of state labels across characters (Lewis, 2001).

In molecular datasets, the state labels unambiguously identify a biochemical structure (‘A’ refers to adenine, ‘C’ to cytosine, and so on), so estimating the frequency of a certain state in a dataset is biologically meaningful. In contrast, state labels in morphological datasets can usually be swapped without any loss of meaning, thus frequencies of ‘0’s and ‘1’s in a dataset are biologically meaningless. Even if meaningful, non-arbitrary state labels are applied to every character, such as ‘0’ = ‘absence’ and ‘1’ = ‘presence’ (Klopfstein et al., 2015), or ‘0’=‘ancestral’ and ‘1’=‘derived’ (Pyron, 2017), those states refer to completely different morphological structures across characters, so modeling their frequencies across a whole dataset is difficult to justify. Because of this arbitrariness of state labels for characters, and lack of common meaning of state labels among characters, a correct phylogenetic model must be invariant to the chosen state labels.

A possible solution to the conundrum of modeling unequal state frequencies in morphological data has been proposed by Wright et al. (2016) for binary characters. In this model, a symmetric beta distribution is discretized into an odd number of categories (five in Wright et al., 2016). The central category represents equal state frequencies, while the other categories represent progressively more unequal state frequencies (these are mirrored so that ‘0’s and ‘1’s can be arbitrarily swapped without resulting in different likelihoods). A shape parameter dictates how unequal state frequencies can be. Different state frequencies for each character are integrated over the likelihood calculation, similarly to how different rates are treated in the discretized gamma model of ACRV.

Applying a variable character acquisition bias correction to a substitution model that allows for unequal state frequencies is not trivial. In fact, the probability of a character being observed as variable depends on its state frequencies. Intuitively, a character with one state that is overwhelmingly more frequent than the other(s) has a very low probability of being observed as variable even when evolving on a tree with long branches. Thus, the tree likelihood conditional on a character being observed as variable is different for different values of the discretized beta distribution from the Wright et al. (2016) model. This means that—exactly as for models with ACRV—there are two different ways to apply the variable character acquisition bias when allowing for unequal state frequencies: a joint acquisition bias approach (jMkv) that conditions the likelihood by state frequency, and a marginal acquisition bias approach (mMkv) that conditions the likelihood over all state frequencies.

While exploring how different acquisition bias approaches impact phylogenetic estimates when allowing for unequal state frequencies—and how that can be combined with ACRV models—is beyond the scope of this paper, we recommend the use of the marginal acquisition bias correction on similar grounds as we did for ACRV models. Namely, marginal acquisition bias better reflects the assumption that characters with more unequal state frequencies are more likely to be observed as invariant—and thus not included in a morphological character matrix—than characters with less unequal state frequencies. Thus, characters with strongly asymmetric state frequencies will be underrepresented in a dataset where all characters are observed to be variable. This fundamentally implies that models with full continuous distributions of base frequencies, or where each character has their own set of independent base frequencies, is not possible as the marginal acquisition bias cannot be computed. While for binary characters we can discretize the base frequency distribution (Wright et al., 2016), this is far more challenging for multi-state characters and requires further research.

### 5.5 Implications for Ancestral State Estimation

Finally, we would like to point out that our findings on modeling variable character acquisition bias might not only be restricted to phylogenetics, but also applicable to ancestral state estimation. Ancestral state estimation of discrete phenotypical characters (not only morphological, but also ecological, physiological, …) is a common analytical approach in macroevolution that falls under the broader umbrella of phylogenetic comparative methods (Pagel, 1994, 1999; Joy et al., 2016; Revell, 2024). The simplest model for ancestral state estimation (the equal-rates model; Schluter et al., 1997; Mooers and Schluter, 1999) is identical to the Mk model of character evolution used in phylogenetics. The only difference lies in the application of the two models, as in ancestral state estimation the phylogeny is usually treated as known with certainty or with some degree of uncertainty, and the character evolution model is applied to one or few focal characters.

To our knowledge, nobody has ever proposed to apply a variable character acquisition bias correction to the likelihood calculation of ancestral character states. We argue that every empirical study with a discrete ancestral state estimation *a priori* selects features that are observed to vary within a taxonomic sample—an ancestral state estimation of a character that is identical among all sampled taxa would not be very informative or interesting after all. Thus, not conditioning the likelihood on the character to be variable would overestimate the evolutionary rate of that character.

The overestimation of the evolutionary rate can have a direct impact on ancestral state estimates, as characters with higher rates (i.e., more labile) are reconstructed with higher uncertainty at internal nodes. While the rate overestimation might be relatively minor (as in the tree length difference between Mk and Mkv models in our simulation study), the effect on reconstructed ancestral states needs to be further explored.

Given the complexity of current models of character evolution used for ancestral state estimation (including varying rates through time and lineages, hidden rates, and correlated evolution; see Revell and Harmon, 2022 for a practical overview), we expect that our discussion on modeling variable character acquisition bias under rate-heterogeneous models will be relevant for further development of phylogenetic comparative methods.

### 5.6 Conclusions

In this paper, we described two distinct approaches to condition tree likelihood on observing variable characters when there is among-character rate variation (ACRV): a joint acquisition bias approach (jMkv model) and a marginal acquisition bias approach (mMkv model). Our results demonstrate that using one or the other acquisition bias correction systematically impacts phylogenetic estimates, including tree length, amount of ACRV, and tree topology. Based on the results of our simulation study and on theoretical arguments, we recommend the use of mMkv + ACRV models for the phylogenetic analysis of morphological datasets. Finally, future development of more complex models of morphological evolution (e.g., character-specific rate heterogeneity, or unequal state frequencies) must consider the impact of different acquisition bias corrections on the likelihood calculation.

## 7 Acknowledgements

We thank xx and yy for comments that helped improve the manuscript. We thank the rest of the Höhna Lab at LMU Munich for helpful discussions.

