## Supplementary Figs. for "On the Mkv Model with Among-Character Rate Variation"

### Supporting Information

[1, 2]Alessio Capobianco [1, 2]Sebastian Höhna
[1] *GeoBio-Center, Ludwig-Maximilians-Universität München, 80333 Munich, Germany*
[2] *Department of Earth and Environmental Sciences, Paleontology & Geobiology, Ludwig-Maximilians-Universität München, 80333 Munich, Germany*

This document is a supplement to the manuscript 'On the Mkv Model with Among-Character Rate Variation'. It consists of two supplementary figures.

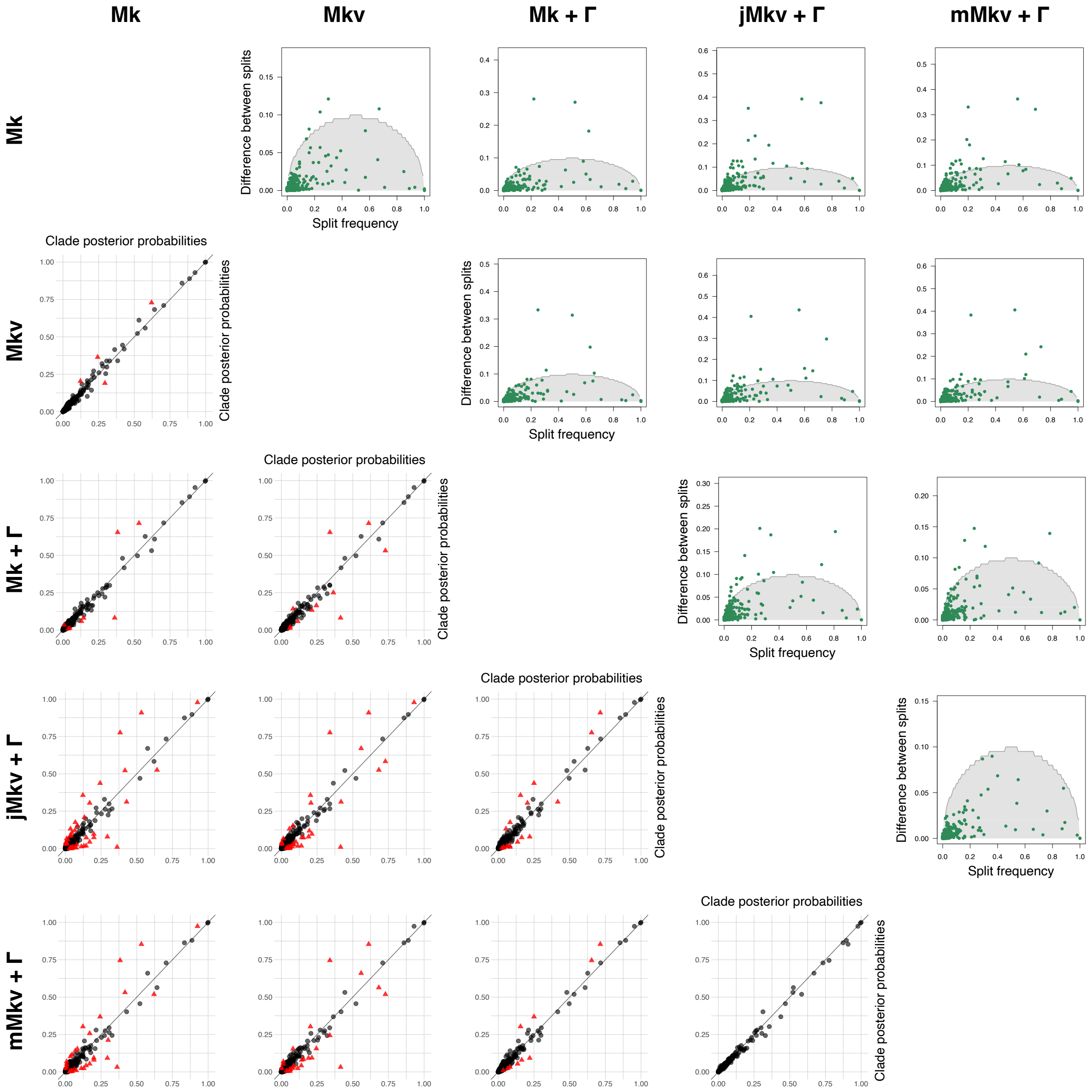


**Figure 1:** Impact of different models of morphological character evolution on tree topology for the Gekkota dataset including both extant and extinct species. Plots below the diagonal are scatter plots showing posterior probabilities of tree splits (or clades) for each pair of models. Each symbol in the scatter plots represents one clade. Circles indicate clades with posterior probabilities not significantly different under the expected difference of split frequencies test. Triangles indicate clades with significantly different posterior probabilities under the same test. Plots above the diagonal show differences between split probabilities for each pair of models, compared to the expected difference of split frequencies corresponding to ESS=200. Each tree split is a dot. When a tree split is above the semicircle, it failed the expected difference of split frequencies test, meaning that it was sampled with a significantly different probability between the two compared models.

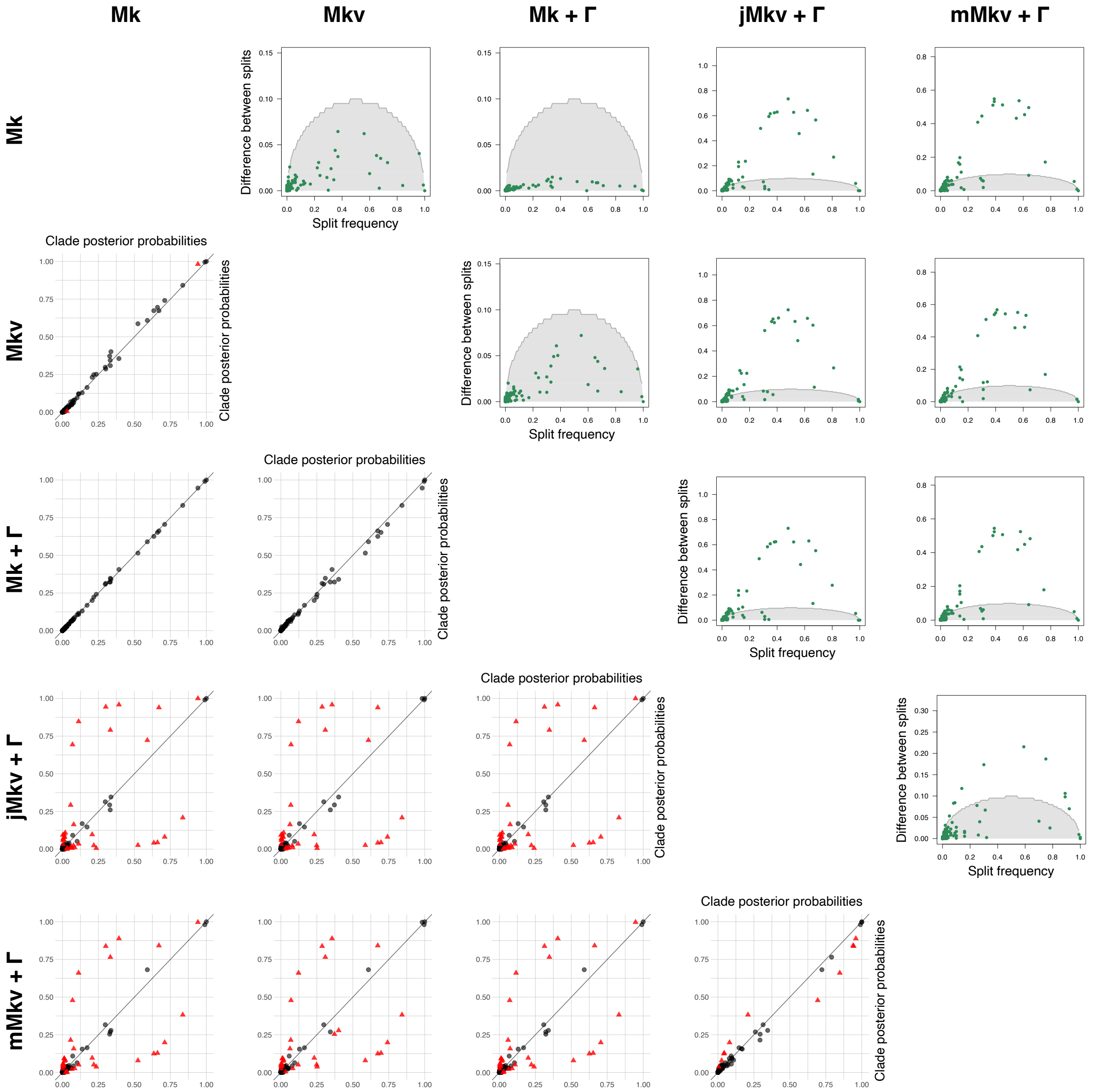


**Figure 2:** Impact of different models of morphological character evolution on tree topology for the Gekkota dataset including only extant species. Plots below the diagonal are scatter plots showing posterior probabilities of tree splits (or clades) for each pair of models. Each symbol in the scatter plots represents one clade. Circles indicate clades with posterior probabilities not significantly different under the expected difference of split frequencies test. Triangles indicate clades with significantly different posterior probabilities under the same test. Plots above the diagonal show differences between split probabilities for each pair of models, compared to the expected difference of split frequencies corresponding to ESS=200. Each tree split is a dot. When a tree split is above the semicircle, it failed the expected difference of split frequencies test, meaning that it was sampled with a significantly different probability between the two compared models.
